# Accelerated matrix-vector multiplications for matrices involving genotype covariates with applications in genomic prediction

**DOI:** 10.1101/2023.07.06.547949

**Authors:** Alexander Freudenberg, Jeremie Vandenplas, Martin Schlather, Torsten Pook, Ross Evans, Jan ten Napel

**Author notes:** Corresponding author, Chair of Applied Stochastics, University of Mannheim, B 6, 26, Mannheim, Germany.

## Abstract

In the last decade, a number of methods have been suggested to deal with large amounts of genetic data in genomic predictions. Yet, steadily growing population sizes and the suboptimal use of computational resources are pushing the practical application of these approaches to their limits. As an extension to the C/CUDA library *miraculix*, we have developed tailored solutions for the computation of genotype matrix multiplications which is a critical bottleneck in the empirical evaluation of many statistical models. We demonstrate the benefits of our solutions at the example of single-step models which make repeated use of this kind of multiplication. Targeting modern Nvidia^®^ GPUs as well as a broad range of CPU architectures, our implementation significantly reduces the time required for the estimation of breeding values in large population sizes. *miraculix* is released under the Apache 2.0 license and is freely available at https://github.com/alexfreudenberg/miraculix.

## 1 Introduction

Over the past 15 years, the incorporation of genomic information has become essential for ensuring progress in breeding (Schaeffer, 2006). A routine task in animal breeding is the estimation of breeding values within a population, that is, the estimation of the average additive effects of the alleles that an individual passes on to its offspring. Though breeding values are estimated most accurately through the use genomic data, it is usually too costly to genotype a whole population. This is particularly true when analyzing large populations, like national dairy evaluations with millions of animals (Misztal et al., 2022). This circumstance has sparked a debate on how to combine pedigree information of ungenotyped animals with Single Nucleotide Polymorphism (SNP) data of genotyped animals to analyze phenotypic records.

Due to its desirable statistical properties, the single-step genomic BLUP (ss-GBLUP) model (Legarra et al., 2009; Christensen and Lund, 2010) has gained popularity in animal breeding, combining both the genomic relationship matrix (GRM) **G** and the pedigree-based relationship matrix **A** into a generalized relationship matrix **H**. Large population sizes used in animal breeding programs, ranging from tens of thousands to millions of individuals, have motivated research in computational strategies to solve the associated mixed model equations (MME) efficiently through the use of high-performance computing (HPC) techniques. One of the proposed approaches is the algorithm for Proven and Young Animals (APY), which approximates the inverse of the GRM, **G**^*−*1^, through genomic recursion on a subset of core animals (Misztal et al., 2014). As an alternative concept, the original ssGBLUP model was reformulated to, first, allow the use of well-established numerical software (Legarra and Ducrocq, 2012) and, second, avoid the explicit construction and inversion of the GRM **G** using the Woodbury decomposition. The resulting models were coined single-step GT(A)BLUP models (Mäntysaari et al., 2017; Mäntysaari et al., 2020). Additionally, single-step SNP BLUP (ssSNPBLUP) models were proposed to estimate SNP effects directly, similarly avoiding **G** and its inverse (Liu et al., 2014; Fernando et al., 2014; Taskinen et al., 2017).

Considering the computational aspects of these approaches, the use of highly-optimized sparse matrix operations has been established, thanks to the sparse characteristics of the pedigree-based relationship matrix. Additionally, established iterative-solver algorithms (e.g., the preconditioned conjugate gradient (PCG)) can be employed in the resulting equation systems (Strandén and Lidauer, 1999; Misztal et al., 2009) to avoid explicit construction of the full coefficient matrix.

Nevertheless, the computational load of the remaining mathematical operations and slow PCG convergence have been somewhat prohibitive to the application of ssGTABLUP and ssSNPBLUP models in ultra-large-scale settings. To mitigate this issue, a number of numerical advances have been proposed to improve convergence speed (Vandenplas et al., 2018). So far, however, improvements in accelerating the involved matrix arithmetics have been limited to the application of shared-memory parallel libraries, such as the Intel^®^ Math Kernel Library (MKL) and PARDISO (Alappat et al., 2020), and unpacking the compressed SNP matrix **M** into the CPU cache for the matrix-matrix product in the PCG iteration (Misztal et al., 2009; Vandenplas et al., 2020). Bit-level algorithms have been employed by, for instance, the popular PLINK software (Chang et al., 2015) in the efficient implementation of genome-wide association studies (GWAS), which rely on similar genotype matrix operations. Yet, these routines are not accessible for use in single-step BLUP evaluations. The software PLINK also implements BLAS-based routines for the calculation of **G** on a Nvidia^®^ GPU in its version 2.0. However, this functionality works on uncompressed SNP data stored in single-precision floating-point values, thereby significantly limiting possible problem sizes. Other authors have used GPUs to accelerate model training for machine learning algorithms in genomic selection (Xu et al., 2021). In parallel to this work, efficient approaches for the multiplication of matrices of mixed input data types have been suggested for use in transformer machine learning models, in particular with sub-byte integer data formats (Kim et al., 2022).

In this study, we present tailored algorithms for the multiplication of a compressed SNP matrix by a matrix of small width stored in floating-point format for CPUs and Nvidia^®^ GPUs. Our CPU code is optimized for all major instruction set architectures. To take advantage of the instruction-level parallelism capabilities of modern CPUs, our implementation uses Single Instruction-stream Multiple Data-stream (SIMD) operations explicitly (Tanenbaum, 2016). Extending the Nvidia^®^ CUTLASS library (Thakkar et al., 2023), our GPU approach benefits from fast tile iterators in the data movement during the matrix-matrix multiplication.

We demonstrate how these advances can drastically reduce the computing times of genotype matrix multiplications on CPUs and GPUs compared to double-precision matrix multiplication routines provided by the Intel^®^ MKL, while simultaneously reducing memory requirements. We provide scripts for reproducing our results in the GitHub repository. Additionally, using genomic data provided by the Irish Cattle Breeding Federation and the genetic evaluation software MiXBLUP (ten Napel et al., 2021), we show how our novel approaches bring down total run times in solving single-step evaluations by up to 62%, thereby paving the way to include even larger population sizes in genomic evaluations.

We provide our implementation as part of the C/CUDA software library *miraculix* (https://github.com/alexfreudenberg/miraculix). Interfaces to call the library from higher-level languages such as Fortran, R and Julia are provided. Through the modular structure of the *miraculix* library, which also supplies functions for the calculation of **G**, the code should be easily modifiable by researchers and practitioners interested in accelerating computations in other BLUP models (Meuwissen et al., 2001; VanRaden, 2008) or other genomic analyses.

## 2 Methods

The efficient multiplication of the genotype matrix by a double-precision matrix plays an important role in many genomic analyses. In this section, we explain how this multiplication can be decomposed to reduce the computational costs involved. Then, we propose novel techniques for the multiplication of a compressed SNP matrix with a double-precision matrix on CPUs and GPUs. We illustrate the role of an efficient genotype matrix multiplication at the example of single-step models. Lastly, we describe the methodology which we used for evaluating our approaches.

### 2.1 Computational bottlenecks

We consider the commonly encountered operation of multiplying the centered SNP matrix **Z**, or its transpose **Z**^*′*^, by a matrix of low width. Here, the matrix **Z** can be computed as

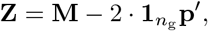

where 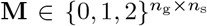 denotes the uncentered genotype matrix containing the *n*_s_ SNP genotypes (coded as 0 for one homozygous genotype, 1 for the heterozygous genotype, or 2 for the alternate homozygous genotype) of *n*_g_ genotyped animals. Furthermore, **p** denotes the vector of allele frequencies and the subtraction of 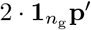 centers the columns of **M**.

Considering an arbitrary matrix 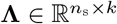, we first note that the multiplication of **Z** with **Λ** can be reformulated into

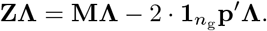

Since subtracting the matrix 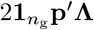 consists of a vector-matrix multiplication and subsequent additions of matrices of rank one, it is computationally cheap and can be achieved with BLAS Level-1 operations. The multiplication of the transposed matrix 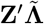 with an arbitrary matrix 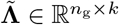 is decomposed similarly into

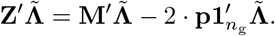

Here, we use the distinction between **Λ** and 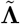 to emphasize that **Z** and **Z**^*′*^ cannot be multiplied with the same matrix because they have different dimensions. Due to the low cost of SNP genotyping, the matrix **M** can have extremely large dimensions, capturing the genomic information of millions of animals. Furthermore, whereas **Λ** is a regular matrix of double-precision, the SNP matrix is usually stored in a compressed format to save memory. For instance, the PLINK 1 binary file format stores the genotypes of four individuals in eight bits (corresponding to one byte), utilizing that one entry of **M** only requires 2 bits of storage. This compressed data format prevents naive calls to BLAS routines, and decompressing it explicitly is inefficient and increases memory requirements. Only recently, the problem of matrix multiplication of mixed input data types has gained attention (Kim et al., 2022) and no off-the-shelf solution exists for compressed 2-bit integer data types.

In general, algorithms for genotype matrix multiplications that operate on compressed data can be expected to be more efficient due to the better utilization of memory movements. In cases where the multiplication **ZΛ** needs to be evaluated repeatedly for varying **Λ**, an optional conversion of the uncentered SNP matrix **M** to a different storage format is comparatively cheap. Therefore, switching the storage format has the potential to yield efficiency gains.

### 2.2 Acceleration of the genotype matrix-matrix multiplication

#### 2.2.1 Previous approach

Vandenplas et al. (2020) proposed a decompress-on-the-fly approach, consisting of unpacking tiles of submatrices of **M** small enough to store the result in the cache and perform matrix multiplication on these tiles. Briefly, modern computer architecture implements different levels of cache memory (commonly, L1, L2 and L3) to reduce access times to repeatedly processed data. While infrequently used data can be stored in the random access memory (RAM) or even on the disk, accessing it over the memory bus combined with the lower clock cycles of the RAM compared to the CPU dramatically slows down the execution of the program. While low levels of cache close to the core allow faster memory reads, they come with lower capacity (Tanenbaum, 2016). Hence, efficient code which processes large amounts of data strives to reduce data movement along the memory hierarchy and utilizes the fast access times of low-level caches. In the aforementioned local decompression approach by Vandenplas et al. (2020), small submatrices of SNP data in PLINK 1 binary format are sequentially converted to double-precision floating-point values. Since these small submatrices are used repeatedly in a loop, they should not be evicted from the L1 cache. Therefore, the unpacked SNP data is readily at hand when new tiles of **Λ** are loaded and can be multiplied without additional conversion operations.

#### 2.2.2 The *5codes* algorithm for CPUs

Building on the idea of keeping frequently used data close to the core, our novel approach for CPU computations aims to reduce data streams of **Λ** through the cache hierarchy by fully avoiding decompression. To explain this approach, we assume that there are no missing values at this point. Instead, they are coded as zero and their effect is taken into account when subtracting 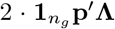 or 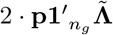 later. Since the number of missing values is usually not substantial, this correction comes at a low cost.

We view the problem of storing compressed SNP data through the lens of combinatorics: Since there are only three possible states (0, 1 and 2) for a SNP, any vector **m** ∈ {0, 1, 2} ^5^ of five contiguous SNPs can assume only one of 3^5^ = 243 values. Assuming there are no missing values, this format requires only 80% of the memory required for storing SNP values in 2 bits. Hence, each realized vector **m** can be stored in one 8-bit unsigned integer while preserving the order of the SNPs. During preprocessing, we convert the input data to this compressed format, which we coined *5codes*. At multiplication time, we treat the columns of **Λ** separately and load a vector *λ* ∈ ℝ^5^ of five entries in a column of **Λ**, which is stored in double precision. Subsequently, we compute all possible results of the scalar product

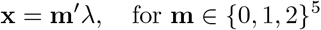

and store them in a hash table. Then, we iterate over the five columns of **M** under consideration and look up the values of **x** depending on the realization of **m**. We provide pseudo-code for one iteration of this approach in Algorithm 1. The multiplication 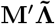 is achieved analogously.

It is worth noting that at least 16 of these tables fit into the L1 cache because each hash table holds 3^5^ = 243 double-precision values of 64 bits and the L1 cache in modern CPU architectures typically holds between 32KB and 96KB. Since the hash table of the different values of **x** only needs to be computed once for every five-row tile of **Λ**, we keep register instructions to a minimum. Furthermore, compressed matrix values are loaded into integer registers whereas real values are stored in SIMD registers. The implementation aims to optimize load operations and store operations. For instance, if SIMD registers of 256-bit width are available, one entry of the hash table contains a vector of four double-precision floating-point values.

The computation is parallelized among the available processor cores by splitting **M** and **Λ** into chunks along the column axis and row axis respectively. Finally, the results of each thread are united by a single reduction operation at the end that computes the sum of all individual results.

As discussed above, genotype centering is not a bottleneck due to its low complexity. Thanks to the structure of the *5codes* encoding, the centering can actually be included in the hash table for the operation **ZΛ**, meaning that instead of holding the possible values of **m**^*′*^*λ*, the hash table can store the centered values

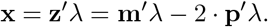

Subtracting 2 · **p**^*′*^*λ* at this point further reduces the number of memory accesses and decreases numerical accumulation errors.

#### 2.2.3 SNP matrix multiplication with mixed data types on GPUs

To make use of the powerful HPC capabilities of modern Nvidia^®^ GPUs, we also implement a matrix multiplication routine in CUDA, in which we extend the CUTLASS library (Thakkar et al., 2023) in its version 2.10. CUTLASS is a C++ template library for high-performance matrix operations on Nvidia^®^ GPUs. In contrast to CPU operations, GPU functions need to align parallel operations both within thread blocks as well as within warps of threads (Sanders and Kandrot, 2010). CUTLASS assists this task by providing a framework that allows software solutions to target only a subset of levels in this hierarchy. In our case, we implement a template specialization for the Single Instruction Multiple Threads (SIMT) subsection within warp-level API in CUTLASS. Since computing times for our GPU approach are mostly driven by data movements instead of algebraic operations, we do not make use of the *5codes* algorithm in this implementation but distribute scalar products of four-dimensional vectors to the cores of the GPU. Therefore, we keep the established PLINK 8-bit-sized format for storing a four-dimensional vector corresponding to four SNPs, where each SNP is coded as either 0 for the homozygous genotype, 1 for a missing genotype, 2 for the heterozygous genotype and 3 for the alternate homozygous genotype. For compatibility with CUTLASS, we introduce a new interleaved data type for double-precision vectors of size four. Furthermore, fitting the genotype matrix multiplication into the CUTLASS framework requires adding a new scalar-product microkernel for this specific combination of data types and adjusting the CUTLASS interfaces upstream accordingly. The microkernel uses bitmasks to extract the SNP values from the compressed storage format and converts them into double-precision floating-point values for immediate multiplication afterward. Through the highly efficient memory access iterators in CUTLASS, we are able to move data quickly from the device memory to the shared memory to the cores and back. Furthermore, in order to reduce memory allocations and data movement between the host and the device, we preallocate memory for the matrix **Λ** and transfer data objects which are required in every PCG iteration (that is, **M** and **p**) only once at start-up time. To our knowledge, this is the first implementation of matrix multiplication of 2-bit integers with double-precision floating-point values, which we designed in parallel to the recent work of Kim et al. (2022) who extended the CUTLASS library to include, among others, matrix multiplication of 4-bit integers with half-precision floating-point values.

##### Algorithm 1

Pseudo-code for the *5codes* algorithm for the multiplication of five columns of SNPs **M** with a vector *λ* of length 5. The multiplication of five columns of **M**^*′*^ is analogous. Note that the variable idx in the algorithm is smaller than 243. For a general number of SNPs *n*_s_, five columns of **M** are multiplied at a time.

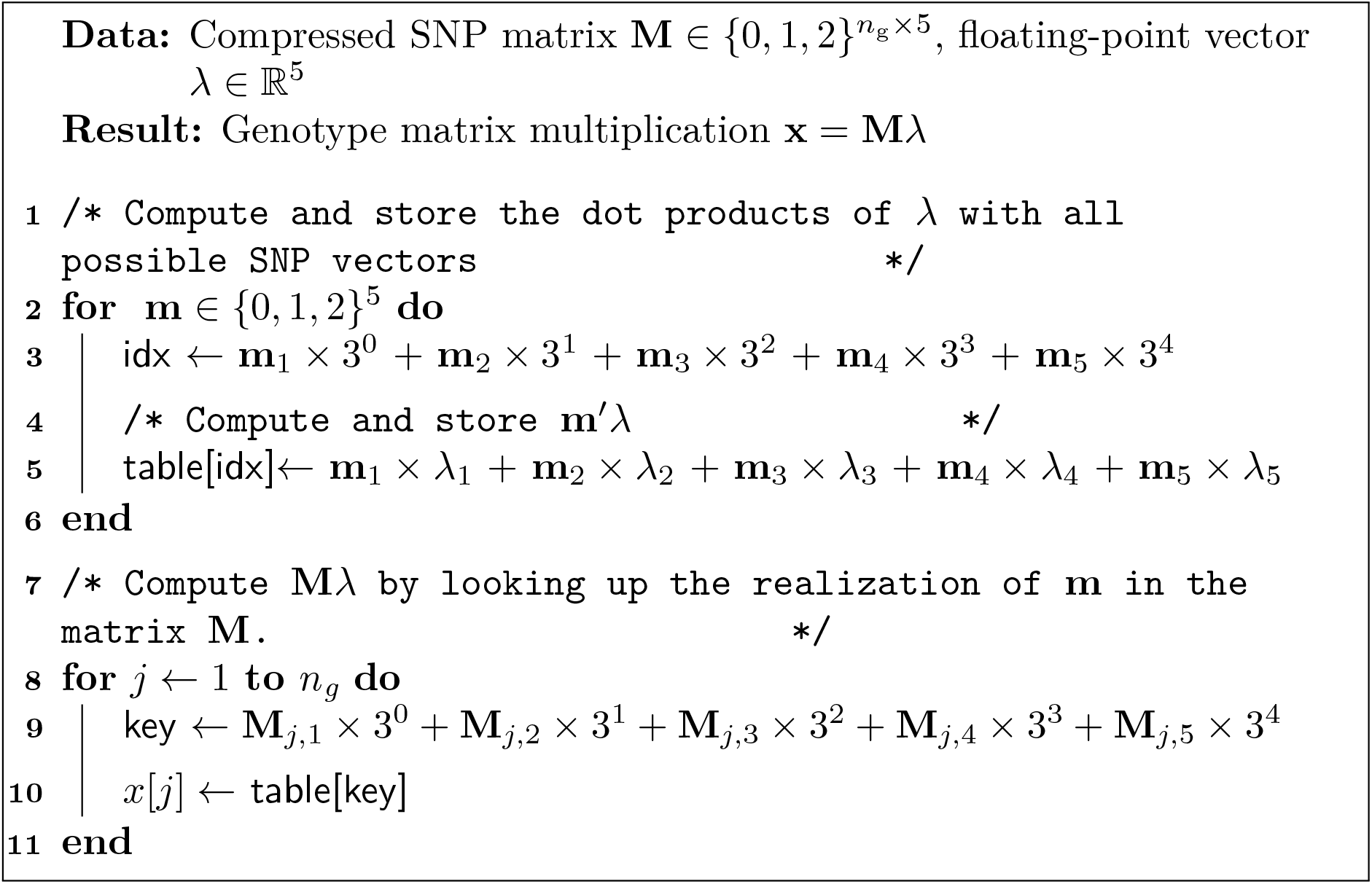

#### 2.2.4 Memory management

Since both the CPU and GPU approaches exploit a compressed storage format for the SNP matrix, the question arises of how to efficiently calculate the transposed matrix product 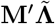 memory-efficient implementation would transpose chunks of **M** of low dimension which are iterated over. For instance, transposing sub-matrices of dimensions 16 by 16 in PLINK 1 binary format would allow distributing the transpose operation in a warp of threads on a GPU. However, thanks to the compressed data storage, we are not memory-bound with the current number of genotyped animals and SNPs, and we, therefore, choose to transpose **M** as a whole during start-up and store it separately to reduce computation time associated with transposition. For instance, the dataset of 2.61m animals with 47k SNP markers we use in this article for testing purposes only requires about 57 gigabytes of RAM for both **M** and **M**^*′*^ combined.

### 2.3 SNP matrix multiplications in single-step models

Single-step models in animal breeding programs commonly comprise hundreds of thousands or even millions of animals. Therefore, the MME for these models need to be solved iteratively in practice, commonly through the PCG algorithm. To this end, each iteration requires multiplying the corresponding coefficient matrix with a candidate vector.

To illustrate the necessity of a fast genotype matrix multiplication, we give an overview of the matrix operations involved in the ssSNPBLUP approach, proposed by Liu et al. (2014), and the ssGTABLUP approach, introduced by Mäntysaari et al. (2017). Both univariate models can be easily extended to multivariate applications. The numerical treatment of these approaches is described in detail by Vandenplas et al. (2023) who found that they have similar computational costs per iteration when applied to large datasets since they require the same matrix computations.

A standard univariate mixed model for ssGBLUP can be written as:

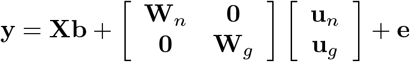

where **y** is the vector of records, **b** is the vector of *n*_fixed_ fixed effects, **u**_*n*_ is the vector of additive genetic effects for the non-genotyped animals, **u**_*g*_ is the vector of additive genetic effects for the genotyped animals, and **e** is the vector of residuals. The matrices **X, W**_*n*_, and **W**_*g*_ are incidence matrices relating records in **y** to the corresponding effects. The random effects vector **u**_*g*_ can be decomposed into **u**_*g*_ = **a**_*g*_ + **Zg**, where **g** is the vector of SNP effects and **a**_*g*_ contains the residual polygenic effects.

Due to the assumed covariance structure, the ssSNPBLUP system of equations proposed by Liu et al. (2014) involves the matrix **Σ**^*−*1^ defined as

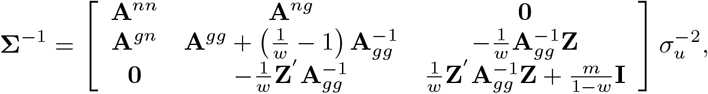

where the scalar 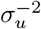 is the inverse of the additive genetic variance, *w* between 0 and 1 is the proportion of variance not explained by SNP markers, called residual polygenic effects, and 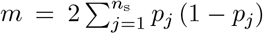 for 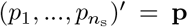. Furthermore, the matrix

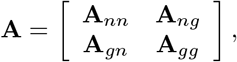

denotes the pedigree-based relationship matrix and

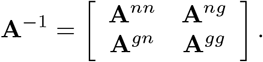

is its inverse, which is sparse (Henderson, 1976). Here, *n* and *g* denote non-genotyped and genotyped animals respectively. The inverse of **A**_*gg*_ can be computed as

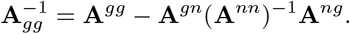

Because the coefficient matrices of the ssSNPBLUP and ssGTABLUP models are too large to be constructed explicitly, it is decomposed into submatrices which are multiplied separately (Vandenplas et al., 2020). Importantly, most of these submatrices (e.g., **A**^*nn*^, **A**^*ng*^, **A**^*gn*^, **A**^*gg*^) in **Σ**^*−*1^ are sparse and can be multiplied by a vector or a matrix at a relatively low cost by using iteration-on-data techniques (Schaeffer and Kennedy, 1986) and sparse matrix operations (Legarra et al., 2009; Vandenplas et al., 2020). Additionally, although the matrix (**A**^*nn*^)^*−*1^ is not sparse in general, equation systems involving (**A**^*nn*^)^*−*1^ can be solved by using forward and backward substitution techniques with the Cholesky factor of **A**^*nn*^, which only needs to be computed once. Therefore, this calculation is also not computationally demanding in practice. In contrast, the multiplication of **Z** by an arbitrary matrix of low width **Λ** has been a computational bottleneck so far.

### 2.4 Evaluation

Since both the *5codes* algorithm and our GPU approach take advantage of modern hardware architectures, it can be expected that they outperform the existing implementations. To quantify the benefits, we used simulated data to benchmark our concepts for genotype data of various sizes. Afterward, considering the ssSNPBLUP and ssGTABLUP systems of equations on a large population of Irish beef and dairy cattle, we evaluated the wall clock times for estimating the breeding values in this population using the PCG implementation in the software MiXBLUP (ten Napel et al., 2021). We compared our novel approaches with the wall clock time required by the current Fortran-based matrix multiplication implementation in MiXBLUP.

#### 2.4.1 Simulated data

For benchmarking our two novel approaches, we simulated genotype data of various dimensions using the software suite PLINK (Chang et al., 2015). Mimicking the population sizes of many breeding programs in practice, we generated genotype data of three distinct animal populations with a varying number of individuals: a small population with 102k animals, a medium population with 751k animals and a large population with 3.1m animals. For each population, 50,241 SNPs were simulated, resulting in memory requirements of approx. 1.2 GB in compressed storage format (38.2 GB in double-precision) for the small population, 8.8 GB (281.2 GB) for the medium population, and 36.3 GB (1160.6 GB) for the large population. Furthermore, we simulated a matrix **Λ** of 10 normally distributed traits.

#### 2.4.2 Cattle data

We tested our two novel approaches when integrated into the PCG solver using data from the routine six-trait calving-difficulty evaluation for Irish dairy and beef cattle performed by Irish Cattle Breeding Federation (ICBF; Ireland) in March 2022. We solved the equation systems associated with the ssSNPBLUP and ssGTABLUP models. The single-step genomic evaluations were based on the same multi-trait animal model and variance components as the current official routine breeding value evaluation described in more detail in Evans et al. (2019) and Vandenplas et al. (2023). Briefly, after extraction and editing, the data file included 16.59m data records (across 6 traits), and the pedigree included 26.46m animals. The genotypes of 2.61m animals included 47,006 SNP markers from 29 bovine autosomes, with a minor allele frequency greater or equal to 0.01. The genotype data were from a range of 30 different arrays ranging from 3k to 850k SNPs that had been imputed using FImpute (Sargolzaei et al., 2014) to a 50k SNP set based on version 3 of the International Beef and Dairy (IDB) chip. For both single-step approaches, the genotype matrix was centered using observed allele frequencies and the proportion of residual polygenic effects was set to *w* = 0.20.

## 3 Results

### 3.1 Benchmarks

In Figure 1, we assess the performance of consecutively multiplying **ZΛ** and 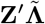, as required in each iteration of a PCG solver. We compare our implementation of the *5codes* algorithm and the GPU implementation with two alternative solutions: 1) The decompress-on-the-fly approach implemented in Fortran and proposed by Vandenplas et al. (2020), which we will call “original solution” below, and 2) a single call to the double-precision matrix multiplication function (dgemm) which is part of all major BLAS libraries. As for the latter, we first needed to inflate the full SNP matrix to double-precision floating-point values and store both the SNP matrix as well as its transpose in memory. We use the BLAS library included in the Intel^®^ Math Kernel Library (MKL). For compilation, we used the Intel^®^ compiler in its version 2021.4.0 and employed compilation options to natively optimize our code to the available hardware. We ran our CPU benchmarks on a single AMD^®^ Milan EPYC 7513 (2.6 GHz) CPU and our GPU implementation on an Nvidia^®^ H100. Since our implementation of the *5codes* algorithm did not display considerable scalability advances beyond 20 cores (see Table 1) and the software MiXBLUP recommends a similar number of cores for parallelizing computations, we refrain from testing it on more cores.

**Table 1:** Computation wall clock times in seconds of the GPU implementation and the CPU algorithm *5codes* for the multiplication of the genotype matrix **Z** and its transposed **Z**^*′*^ with simulated matrices **Λ** and 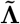. CPU operations were performed with 2 TB of RAM on the Intel^®^ Xeon Gold 6230 (2.1 GHz) and the AMD^®^ EPYC 7513 (2.6 GHz) on 1 core, 10 cores and 20 cores. The V100 GPUs were equipped with 32 GB of device memory, while the A100 and H100 models had a capacity of 80 GB. Dashes (-) indicate out-of-memory events. Results are the average of 10 repeated calculations.

| Population size | Operation | Nvidia<sup>®</sup> |  |  | Intel<sup>®</sup> |  |  | AMD<sup>®</sup> |  |  |
| --- | --- | --- | --- | --- | --- | --- | --- | --- | --- | --- |
|  |  | V100 | A100 | H100 | 1c | 10c | 20c | 1c | 10c | 20c |
| Small | $\mathbf{Z}\mathbf{\Lambda}$ | 0.10 | 0.06 | 0.05 | 2.24 | 0.36 | 0.35 | 1.71 | 0.38 | 0.41 |
| | $\mathbf{Z}'\tilde{\mathbf{\Lambda}}$ | 0.10 | 0.06 | 0.05 | 2.29 | 0.37 | 0.29 | 1.75 | 0.30 | 0.28 |
| Medium | $\mathbf{Z}\mathbf{\Lambda}$ | 0.69 | 0.45 | 0.35 | 16.03 | 2.61 | 2.51 | 12.53 | 2.96 | 2.94 |
| | $\mathbf{Z}'\tilde{\mathbf{\Lambda}}$ | 0.70 | 0.47 | 0.36 | 16.81 | 2.50 | 1.81 | 13.00 | 2.08 | 1.33 |
| Large | $\mathbf{Z}\mathbf{\Lambda}$ | - | 1.87 | 1.42 | 76.42 | 12.31 | 12.24 | 51.72 | 10.99 | 11.03 |
| | $\mathbf{Z}'\tilde{\mathbf{\Lambda}}$ | - | 1.94 | 1.47 | 83.44 | 12.73 | 10.04 | 53.68 | 8.36 | 5.55 |

**Figure 1:**
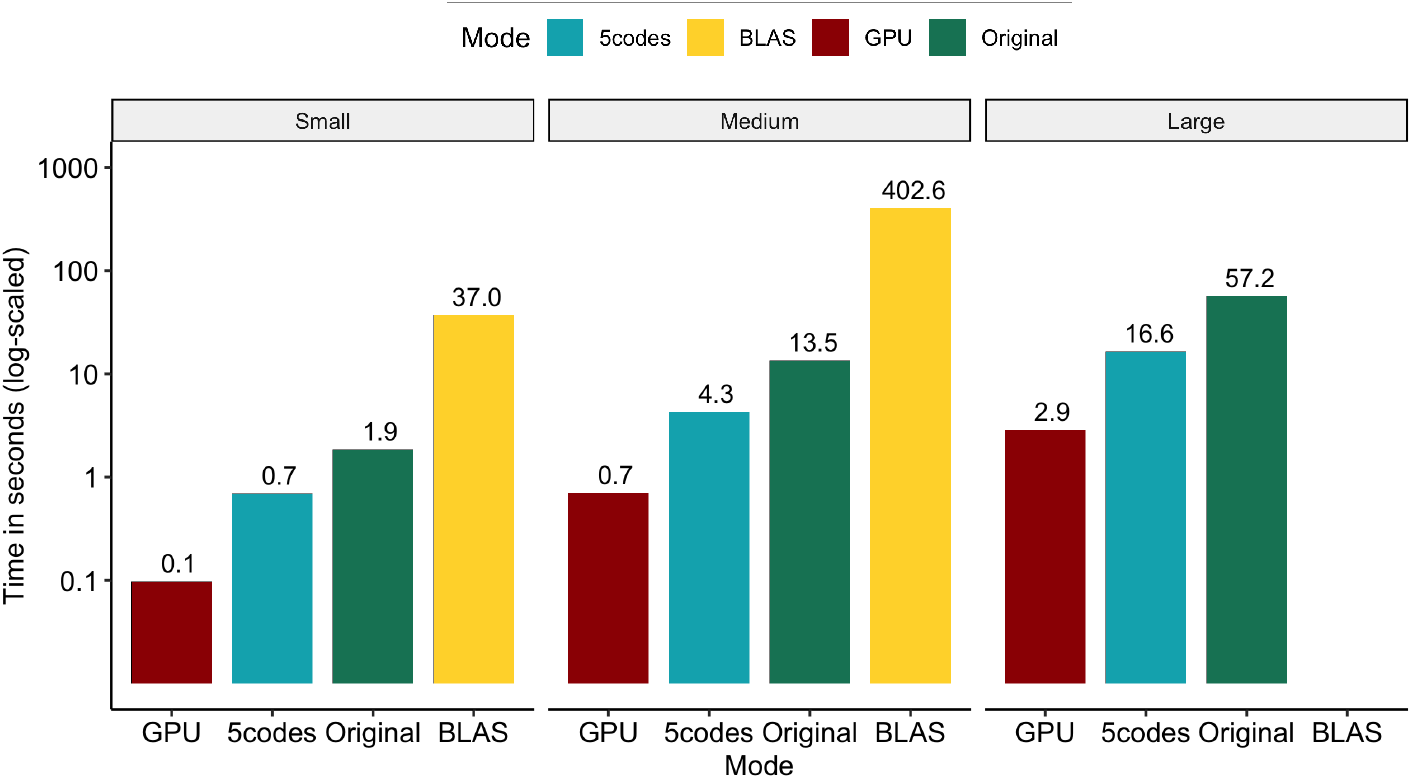
Total wall clock time for consecutively calculating **ZΛ** and 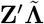 on a system with a dual-socket AMD^®^ Milan EPYC 7513 (2.6 GHz) processor, 2 TB of RAM and a single Nvidia^®^ H100 GPU. Annotations show formatted computing times in seconds. CPU calculations used 20 dedicated cores. No test was performed for the BLAS routine with the large population size, as this would have required ca. 2.27 TB of RAM. Results are the average of 10 repeated calculations.

Evaluating our two novel approaches against a crude call to the BLAS-based double-precision matrix multiplication function shows that computing times can be reduced by more than 99.7% (98.1%) for the small and medium populations on the GPU (CPU). Unfortunately, the genotype data of the large population in double precision required approx. 2.27 TB of RAM, which could not be met in our hardware setup, hindering us from benchmarking the BLAS library on this dataset. Similarly, the CPU-based original solution for multiplying compressed SNP matrices with **Λ** took about 19 times longer on all population sizes compared to the GPU implementation and three times longer compared to the *5codes* algorithm.

To mitigate the impact of hardware-specific optimizations which might limit the reproducibility of our results, we further test our CPU code on Intel^®^ and AMD^®^ processors (Intel^®^ Xeon Gold 6230 and AMD^®^ Milan EPYC 7513) and our GPU code on three generations of Nvidia^®^ datacenter GPUs (Volta V100, Ampere A100 and Hopper H100). Results displayed in Table 1 are the average computing times of 10 replicates.

As for the GPU implementation, our findings suggest that reductions in computing times of approx. 25% (50%) can be achieved when using the recently introduced H100 compared to the A100 (V100). More importantly, the V100 ships with 16 GB or 32 GB of device RAM, limiting the potential problem sizes in our application. Yet, this issue could be mitigated by using multiple GPUs. Since both the transposed and the untransposed matrix are stored on the device, computing times for both multiplications do not differ substantially.

Despite using the Intel^®^ C/C++ Classic Compiler with Intel-specific optimizations, the *5codes* algorithm performs slightly better on the more powerful AMD^®^ chip than on the Xeon Gold 6230. We observe that our implementation scales reasonably well to 10 cores, decreasing computing times by at least a factor of 5. However, increasing the number of cores to 20 only yields mild performance improvements and even comes with a penalty in some cases (see Table 1).

Overall, our evaluations suggest that the GPU implementation significantly outperforms the *5codes* implementation, though practical applications should evaluate the costs and benefits of adding a GPU to their hardware setup based on their respective compute time restraints.

### 3.2 Impact on single-step genomic evaluations

We solved the equation systems associated with the ssSNPBLUP and ssGTABLUP models with the program hpblup, a PCG-based solver used by the software MiXBLUP 3.1 (ten Napel et al., 2021), which links against the *miraculix* library and toggles the use of our two novel implementations through an option. Experiments were performed with 180 GB of RAM on a single AMD^®^ EPYC 7513 CPU for the *5codes* algorithm and a single Nvidia^®^ A100 for the GPU implementation. For the CPU tests, we used 15 dedicated cores, as this is the recommended setting in MiXBLUP. The PCG was iterated until a square relative residual below 10^*−*13^ was achieved. To put our observed computing times into perspective, we also solved both models with the current approach for multiplying genotype matrices implemented in MiXBLUP 3.1 (which we will call “current” below).

Results are displayed in Table 2. Due to the different nature of the effects estimated, the computing times should be evaluated separately for the two models. For the ssSNPBLUP equation system, we observe that the average wall clock time is reduced by approx. 72% (11%) for the multiplication **ZΛ** and by approx. 95% (76%) for 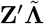 on GPUs (CPUs) respectively. When solving the ssGTABLUP equation system, the average time for multiplying **ZΛ** was slightly higher in the *5codes* algorithm. Yet, both *5codes* and the GPU implementation significantly decreased the time for computing 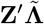.

**Table 2:**
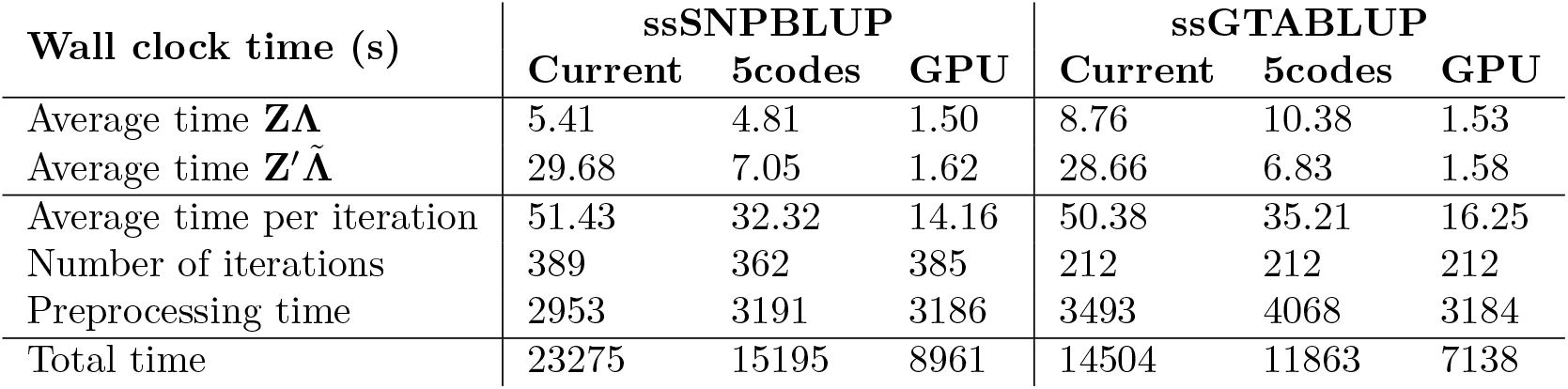
Computation wall clock times of single-step genomic models on ICBF cattle data in seconds. The SNP matrix **Z** and its transposed **Z**^*′*^ are multiplied by matrices **Λ** and 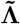 respectively to compute the candidate matrix in the PCG. All computations were performed on one AMD Milan EPYC 7513 CPU with 15 dedicated cores. GPU computations were performed on a single Nvidia^®^ A100 GPU with 80 GB device memory. Both models were trained to a relative error below 10^*−*13^. The total time contains preprocessing (I/O operations and set-up of the preconditioner matrix), solving of the MME, and postprocessing (mainly I/O operations).

As the genotype matrix multiplication constitutes a significant portion of the time per iteration in the PCG solver, the *5codes* implementation reduced the time for solving the ssSNPBLUP model from approx. 6.47 hours to 4.22 hours on the CPU. Similarly, the wall clock time for solving the ssGTABLUP model was reduced from 4.03 hours to 3.30 hours. Outperforming these results, the GPU approach only required 2.49 hours for the ssSNPBLUP model and 1.98 hours for the ssGTABLUP model. A similar number of iterations was observed for solving all models, though the *5codes* algorithm required 27 iterations less for solving the MME associated with the ssSNPBLUP model, which might be explained by its inherently higher precision.

## 4 Discussion

We have presented novel approaches for multiplying centered genotype matrices **M** by a continuously-scaled matrix **Λ** which are applicable both on CPUs as well as on modern GPUs. Useful applications include single-step genomic models that are used to compute breeding value estimates when only a subset of animals are genotyped and/or phenotyped. Yet, similar computational operations are employed in other fields of modern genetics. For instance, genome-wide association studies could benefit from a fast genotype matrix multiplication at various computational bottlenecks: the multiplication of the SNP matrix by a phenotype vector is an essential part of the calculation of genotype-phenotype correlations (Yang et al., 2011). Additionally, many genome-wide association studies use the results of a principal component analysis (PCA) of **G** for population stratification (Price et al., 2006; Meuwissen et al., 2017; Ødegård et al., 2018) and hence are to gain from a fast genotype matrix multiplication as well.

Through our optimized algorithms we were able to achieve a speed-up of critical operations by a factor of up to 3 compared to the methodology by Vandenplas et al. (2020) using CPUs, and a factor of up to 20 using GPUs. Thanks to this acceleration, we have shown how our software library can be used by researchers and practitioners to estimate breeding values in a population of 26.46m animals, 2.61m of which were genotyped, in a reasonable time of approx. 2 hours.

Nevertheless, the growth of breeding populations as well as the steadily falling costs of genotyping will result in genomic datasets of ever-growing size. Therefore, there are several avenues for further research to utilize computing resources even more efficiently.

First, as indicated in Section 2, system memory requirements might be reduced by a factor of approximately two by transposing the compressed genotype matrix on-the-fly during matrix multiplication instead of storing the transpose explicitly. With Nvidia^®^ GPUs currently limited to at most 94 GB device memory and most compute set-ups limited to hundreds of gigabytes of RAM, this improvement would extend the dimensions of possible problem sizes addressable with our proposed matrix-multiplication microkernel.

Second, though our GPU implementation uses highly efficient data access iterators provided by the CUTLASS library, a further reduction in computing time might be achieved by using warp-level-coordinated matrix operations, which have been added as hardware instructions on the latest generations of GPUs (see, e.g., the CUTLASS documentation for how this has been tackled in general tensor-tensor operations (Thakkar et al., 2023)). Additionally, we have seen that our *5codes* implementation suffers from scalability issues when extending the number of cores.

Finally, it should be noted that in the evaluation of single-step models, the preprocessing time required by the PCG solver (e.g. to set up the preconditioner) now constitutes a significant portion of the total computation time. Reducing this contribution holds the potential for additional performance improvements.

Notwithstanding these potential improvements, our software can be used in a variety of computational tasks in genomics to reduce computing times.

## Author Contributions

MS designed and implemented the *5codes* algorithm. AF designed and implemented the GPU solution. AF, JV and MS implemented interfaces to the algorithms. AF and JV wrote and ran analyses for this script. RE preprocessed and curated the ICBF cattle data. TP and JtN provided valuable suggestions for the experiment design and the focus of this study. All authors contributed to the article and approved the submitted version.

## Acknowledgments

The authors thank Associate Editor Dorian Garrick and two reviewers for their review and valuable suggestions to the manuscript.

## Data Availability Statement

The code in the *miraculix* package is released open-source under the Apache 2.0 license and is freely available on GitHub (https://github.com/alexfreudenberg/miraculix). Scripts for simulating the datasets used in this study can be found in the repository. The software *MiXBLUP* and the included solver *hpblup*, integrating the *miraculix* package, is commercially available at https://www.mixblup.eu/.

Individual genotype, pedigree and phenotype data used in this study are managed by the Irish Cattle Breeding Federation and cannot be made publicly available. Reasonable requests for the cattle data can be made to the Irish Cattle Breeding Federation (; website: https://www.icbf.com/). Requests will be considered on a case-by-case basis, subject to approval by the data owner and compliance with any legal and ethical requirements related to data access and use.

## References

Alappat, C., Basermann, A., Bishop, A. R., Fehske, H., Hager, G., Schenk, O., et al. (2020). A recursive algebraic coloring technique for hardware-efficient symmetric sparse matrix-vector multiplication. ACM Trans. Parallel Comput. 7, 1–37. doi:10.1145/3399732

Chang, C. C., Chow, C. C., Tellier, L. C., Vattikuti, S., Purcell, S. M., and Lee, J. J. (2015). Second-generation PLINK: Rising to the challenge of larger and richer datasets. GigaScience 4, s13742–015. doi:10.1186/s13742-015-0047-8. S13742-015-0047-8

Christensen, O. and Lund, M. (2010). Genomic prediction when some animals are not genotyped. Genetics Selection Evolution 42, 2. doi:10.1186/1297-9686-42-2

Evans, R., Cromie, A., and Pabiou, T. (2019). Genetic evaluations for dam-type specific calving performance traits in a multi-breed population. In Book of Abstracts of the 70th Annual Meeting of the European Federation of Animal Science, ed. E. S. et al. 468

Fernando, R. L., Dekkers, J. C., and Garrick, D. J. (2014). A class of Bayesian methods to combine large numbers of genotyped and non-genotyped animals for whole-genome analyses. Genetics Selection Evolution 46, 1–13. doi:10.1186/1297-9686-46-50

Henderson, C. R. (1976). A simple method for computing the inverse of a numerator relationship matrix used in prediction of breeding values. Biometrics 32, 69–83

Kim, Y. J., Henry, R., Fahim, R., and Hassan, H. (2022). Who says elephants can’t run: Bringing large scale MoE models into cloud scale production. In Proceedings of The Third Workshop on Simple and Efficient Natural Language Processing (SustaiNLP), eds. A. Fan, I. Gurevych, Y. Hou, Z. Kozareva, S. Luccioni, N. S. Moosavi, S. Ravi, G. Kim, R. Schwartz, and A. Rücklé (Abu Dhabi, United Arab Emirates (Hybrid): Association for Computational Linguistics), 36–43

Legarra, A., Aguilar, I., and Misztal, I. (2009). A relationship matrix including full pedigree and genomic information. Journal of Dairy Science 92, 4656–4663. doi:10.3168/jds.2009-2061

Legarra, A. and Ducrocq, V. (2012). Computational strategies for national integration of phenotypic, genomic, and pedigree data in a single-step best linear unbiased prediction. Journal of Dairy Science 95, 4629–4645. doi: 10.3168/jds.2011-4982

Liu, Z., Goddard, M. E., Reinhardt, F., and Reents, R. (2014). A single-step genomic model with direct estimation of marker effects. Journal of Dairy Science 97, 5833–5850. doi:10.3168/jds.2014-7924

Mäntysaari, E. A., Evans, R. D., and Strandén, I. (2017). Efficient single-step genomic evaluation for a multibreed beef cattle population having many genotyped animals. Journal of Animal Science 95, 4728–4737. doi:10.2527/jas2017.1912

Meuwissen, T., Indahl, U., and Ødegård, J. (2017). Variable selection models for genomic selection using whole-genome sequence data and singular value decomposition. Genetics Selection Evolution 49, 94. doi:10.1186/s12711-017-0369-3

Meuwissen, T. H. E., Hayes, B., and Goddard, M. E. (2001). Prediction of total genetic value using genome-wide dense marker maps. Genetics 157, 1819–1829. doi:10.1093/genetics/157.4.1819

Misztal, I., Legarra, A., and Aguilar, I. (2009). Computing procedures for genetic evaluation including phenotypic, full pedigree, and genomic information. Journal of Dairy Science 92, 4648–55. doi:10.3168/jds.2009-2064

Misztal, I., Legarra, A., and Aguilar, I. (2014). Using recursion to compute the inverse of the genomic relationship matrix. Journal of Dairy Science 97, 3943–3952. doi:10.3168/jds.2013-7752

Misztal, I., Lourenco, D., Tsuruta, S., Aguilar, I., Masuda, Y., Bermann, M., et al. (2022). 668. How ssGBLUP became suitable for national dairy cattle evaluations. In Proceedings of 12th World Congress on Genetics Applied to Livestock Production (WCGALP), eds. R. F. Veerkamp and Y. de Haas (Wageningen Academic Publishers), 2757–2760. doi:10.3920/978-90-8686-940-4_668

Mäntysaari, E., Koivula, M., and Strandén, I. (2020). Symposium review: Single-step genomic evaluations in dairy cattle. Journal of Dairy Science 103, 5314–5326. doi:10.3168/jds.2019-17754

Ødegård, J., Indahl, U., Strandén, I., and Meuwissen, T. H. (2018). Largescale genomic prediction using singular value decomposition of the genotype matrix. Genetics Selection Evolution 50. doi:10.1186/s12711-018-0373-2

Price, A. L., Patterson, N. J., Plenge, R. M., Weinblatt, M. E., Shadick, N. A., and Reich, D. (2006). Principal components analysis corrects for stratification in genome-wide association studies. Nature Genetics 38, 904–909. doi:10.1038/ng1847

Sanders, J. and Kandrot, E. (2010). CUDA by example: an introduction to general-purpose GPU programming (Addison-Wesley Professional)

Sargolzaei, M., Chesnais, J. P., and Schenkel, F. S. (2014). A new approach for efficient genotype imputation using information from relatives. BMC Genomics 15, 478. doi:10.1186/1471-2164-15-478

Schaeffer, L. (2006). Strategy for applying genome-wide selection in dairy cattle. Journal of Animal Breeding and Genetics 123, 218–223. doi: 10.1111/j.1439-0388.2006.00595.x

Schaeffer, L. and Kennedy, B. (1986). Computing strategies for solving mixed model equations. Journal of Dairy Science 69, 575–579. doi:10.3168/jds.S0022-0302(86)80441-6

Strandén, I. and Lidauer, M. (1999). Solving large mixed linear models using preconditioned conjugate gradient iteration. Journal of Dairy Science 82, 2779–2787. doi:10.3168/jds.S0022-0302(99)75535-9

Tanenbaum, A. S. (2016). Structured computer organization (Pearson Education India)

Taskinen, M., Mäntysaari, E., and Strandén, I. (2017). Single-step SNP-BLUP with on-the-fly imputed genotypes and residual polygenic effects. Genetics Selection Evolution 49. doi:10.1186/s12711-017-0310-9

Ten Napel, J., Vandenplas, J., Lidauer, M. H., Strandén, I., Taskinen, M., Mäntysaari, E. A., et al. (2021). MiXBLUP 3.0.1 manual. Animal Breeding and Genomics, Wageningen University & Research, Wageningen, the Netherlands, v3.0 edn. Last accessed 16th of July 2023

Thakkar, V., Ramani, P., Cecka, C., Shivam, A., Lu, H., Yan, E., et al. (2023). Vandenplas, J., Eding, H., Bosmans, M., and Calus, M. P. (2020). Computational strategies for the preconditioned conjugate gradient method applied to ssSNPBLUP, with an application to a multivariate maternal model. Genetics Selection Evolution 52, 1–10. doi:10.1186/s12711-020-00543-9

Vandenplas, J., Eding, H., Calus, M. P., and Vuik, C. (2018). Deflated preconditioned conjugate gradient method for solving single-step BLUP models efficiently. Genetics Selection Evolution 50, 1–17. doi:10.1186/s12711-018-0429-3

Vandenplas, J., ten Napel, J., Darbaghshahi, S. N., Evans, R., Calus, M. P., Veerkamp, R., et al. (2023). Efficient large-scale single-step evaluations and indirect genomic prediction of genotyped selection candidates. Genetics Selection Evolution 55, 1–17. doi:10.1186/s12711-023-00808-z

VanRaden, P. (2008). Efficient methods to compute genomic predictions. Journal of Dairy Science 91, 4414–4423. doi:https://doi.org/10.3168/jds.2007-0980

Xu, Y., Laurie, J. D., and Wang, X. (2021). CropGBM: An ultra-efficient machine learning toolbox for genomic selection-assisted breeding in crops. In Springer Protocols Handbooks (Springer Protocols Handbooks). 133–150. doi:10.1007/978-1-0716-1526-3_5

Yang, J., Lee, S., Goddard, M., and Visscher, P. (2011). GCTA: a tool for genome-wide complex trait analysis. American Journal of Human Genetics 88, 76–82. doi:10.1016/j.ajhg.2010.11.011

